# YY1-concentration-dependent formation of mechanically distinct DNA condensates through different interaction mechanisms

**DOI:** 10.64898/2026.03.31.715725

**Authors:** Xi Yan, Tsuyoshi Terakawa

## Abstract

The transcription factor Yin Yang 1 (YY1) plays roles in chromatin organization, combining sequence-specific DNA recognition via zinc finger domains with multivalent interactions mediated by intrinsically disordered regions. While YY1 has been implicated in phase separation and enhancer–promoter communication, how its structured and intrinsically disordered regions cooperate to shape DNA–protein assemblies remains unclear. Here, we used single-molecule DNA curtain fluorescence imaging to dissect the molecular basis of YY1-DNA assembly. We found that YY1 induces higher-order assembly in a concentration-dependent manner. At moderately high concentrations, YY1 formed weakly-linked DNA condensates in which dynamic, liquid-like YY1 molecules are scaffolded by relatively immobile, solid-like DNA. At high concentrations, zinc finger–mediated bridging dominates, producing strongly-linked condensates. Distinct domain-deletion mutants selectively impaired either weakly-(soft) or strongly-linked (hard) DNA condensate formation, indicating that these architectures are generated by separate domain-dependent mechanisms. Our findings establish a domain-level framework for DNA condensate formation and highlight how transcription factors can integrate specific and non-specific DNA interactions to control the material state of chromatin and would influence genome regulation.

## INTRODUCTION

Precise regulation of gene expression is fundamental to essential biological processes such as cell growth, differentiation, and development in eukaryotic organisms^1^. Recent advances have revealed that this regulation is governed not only by sequence-specific DNA–protein interactions but also by the dynamic three-dimensional organization of chromatin^2–4^. In particular, the spatial proximity between regulatory elements such as enhancers and promoters is critical for transcriptional control^5–7^, and is often mediated by architectural transcription factors^3,8,9^. These mechanisms, including enhancer–promoter looping and the formation of transcriptional condensates supposedly through liquid–liquid phase separation (LLPS), are now recognized as principles of gene regulation across eukaryotes^10–12^. However, it remains unclear how architectural transcription factors employ their distinct intra-protein domains to influence the physical and dynamic properties of the resulting DNA–protein assemblies. Understanding these mechanisms is relevant, because differences in the material state of these assemblies could profoundly affect the accessibility of transcriptional machinery and thereby modulate gene regulation.

Yin Yang 1 (YY1) has emerged as a unique architectural transcription factor with pivotal roles in organizing chromatin topology (Fig. 1A)^13–15^. YY1 has traditionally been thought to bind to both enhancers and promoters and facilitates their physical interaction through sequence-specific recognition and dimerization^16,17^. Genetic or epigenetic disruption of YY1 binding sites destabilizes enhancer–promoter contacts and diminishes transcriptional output, underscoring its essential role in maintaining higher-order chromatin architecture and promoting gene activation^17^. Genome-wide chromatin interaction analyses further demonstrated that YY1 occupies interacting enhancers and promoters and structurally bridges them through homodimerization, functioning analogously to CTCF in organizing topological domains but acting specifically at enhancer–promoter loops^14^. However, recent acute depletion studies suggest that although YY1 contributes to the establishment of enhancer–promoter contacts, the short-term maintenance of these loops appears largely independent of YY1, implying that additional factors or chromatin-based memory mechanisms sustain loop architecture once formed^19^.

**Figure 1:**
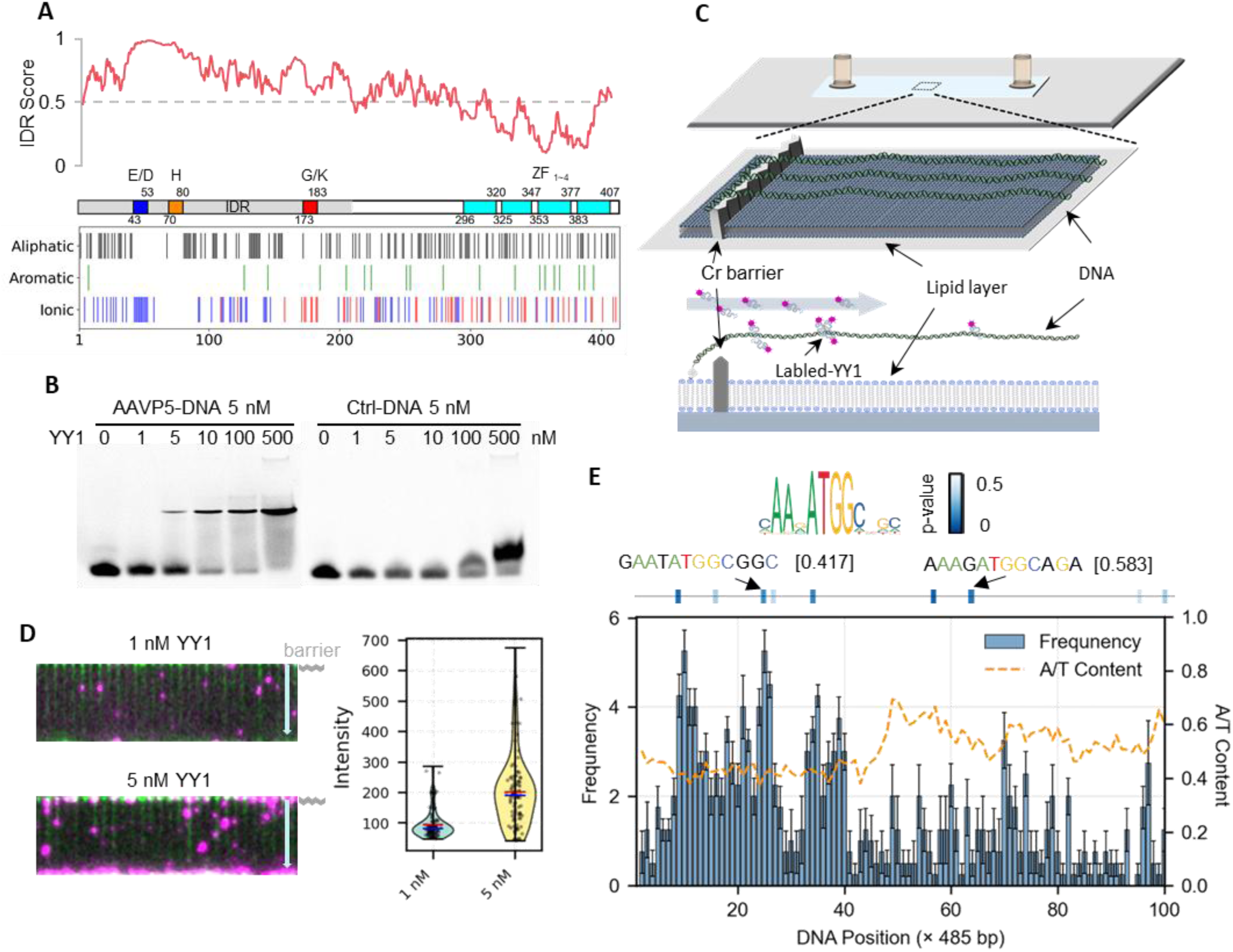
YY1 forms concentration-dependent condensation on DNA through specific and multivalent interactions. **(A)** Intrinsic disorder and domain architecture of human YY1. Top: per-residue disorder probability predicted by IUPred2A. Bottom: schematic of YY1 domains, including the acidic E/D region, histidine-rich region (H), glycine/lysine-rich region (G/K), and four zinc fingers (ZF1–4). The distribution of aliphatic, aromatic, and charged residues is indicated below. **(B)** Electrophoretic mobility shift assay (EMSA). Left: YY1 binds AAVP5 promoter-derived DNA in a concentration-dependent manner. Right: No detectable binding is observed with control DNA. **(C)** DNA curtain assay. λDNA molecules (green) are tethered to a fluid lipid bilayer via biotin–streptavidin interactions, aligned along nanofabricated chromium barriers, and visualized by total internal reflection fluorescence microscopy. Top: schematic illustrating YY1 binding and condensate formation. Bottom: representative kymograph showing DNA (green) and Qdot-labeled YY1 (magenta). **(D)** YY1 condensate formation on DNA. Representative fluorescence images show sparse binding at 1 nM (top) and large condensate formed via multivalent interactions at 5 nM (bottom). Right: violin plots of YY1 fluorescence intensity distributions at 1 nM (blue) and 5 nM (yellow). Red lines indicate the mean; blue lines indicate the median. **(E)** Distribution of YY1 binding along λDNA. The histogram shows the frequency of YY1 binding events (mean ± SEM) per 485 bp bin. The dashed line indicates local A/T content calculated in 10 bp windows. Predicted YY1 consensus motifs are shown as light blue boxes, with a representative motif logo. Examples of motifs with high and low A/T content are indicated, with corresponding p-values in parentheses.

Beyond its sequence-specific binding via zinc finger domains, YY1 exhibits properties of a phase-separating protein in the absence of DNA^20^. Its N-terminal intrinsically disordered regions (IDRs) (Fig. 1A), particularly the histidine-rich region, can drive the formation of dynamic, droplet-like condensates both *in vitro* and in cells^20,21^. These YY1 condensates are enriched at active genomic loci, where they compartmentalize transcriptional coactivators such as p300/CBP, BRD4, and Mediator, as well as RNA polymerase II^22^. Notably, phase separation-deficient YY1 mutants fail to form such condensates and lose their ability to cluster enhancers with promoters, indicating that LLPS is functionally coupled to YY1-mediated gene activation^23^. However, the mechanistic properties of YY1–DNA co-condensates have not been characterized *in vitro*, leaving a gap in our understanding of how YY1 behaves on DNA substrates. Additionally, how YY1’s structured zinc finger domains and disordered low-complexity regions cooperate to organize chromatin and regulate the physical state of DNA–protein assemblies remains poorly understood.

In this study, we set out to dissect how YY1’s structured and disordered domains cooperate to assemble protein–DNA co-condensates and to determine how condensate properties depend on protein concentration. Using single-molecule DNA curtain fluorescence imaging and domain-deletion analysis^24^, we reveal that YY1 engages in modular, concentration-dependent interactions with long double-stranded DNA, giving rise to condensates with distinct physical states. At moderately high concentrations, YY1 forms “soft” multi-DNA condensates, in which dynamic, liquid-like YY1 molecules are scaffolded by relatively immobile, solid-like DNA. These structures arise through cooperation between specific and non-specific DNA binding by the zinc finger domains and non-specific interactions mediated by positively charged subregions within the IDR. In contrast, at high concentrations, zinc finger–mediated bridging dominates, producing mechanically robust “hard” condensates. The ability to selectively disrupt soft or hard multi-DNA condensates through deletion of different YY1 subregions demonstrates that these assemblies are generated by distinct subregion-dependent mechanisms. These results move beyond the conventional view that increasing YY1 concentration simply elevates occupancy at specific binding sites; instead, they support a model in which changes in the physical properties of YY1–DNA co-condensates serve as a regulatory layer that can actively influence downstream transcriptional outcomes.

## RESULT

### Sequence- and context-dependent binding of YY1

To establish a baseline for how YY1 engages with DNA and to assess whether its binding is modulated by protein concentration, we performed electrophoretic mobility shift assays (EMSA)^25^ using increasing concentrations of YY1 (0, 1, 5, 10, 100 and 500 nM) in the presence of either adeno-associated virus P5-derived DNA (AAVP5-DNA; Fig. 1B, left)^13^ or control DNA (Ctrl-DNA; Fig. 1B, right), both at 5 nM. With AAVP5-DNA, we observed that the intensity of the YY1–DNA complex band increased progressively with increasing YY1 concentration. For Ctrl-DNA, which lacks a specific YY1 binding site, no distinct band corresponding to the YY1–DNA complex was observed, even when the YY1 concentration was increased up to 500 nM. These results indicate that YY1 interacts more strongly with AAVP5-derived DNA than with control DNA. For Ctrl-DNA, a slightly upward-shifted band was observed at very high protein concentrations. At extremely high protein concentrations, the overall ionic environment of the sample may be substantially altered. Increased ionic strength and local charge neutralization can reduce electrostatic repulsion along the DNA backbone, potentially affecting its conformation and hydration state, which may account for the appearance of the slightly upward-shifted band observed under these conditions. Even when the protein concentration was increased up to 500 nM using DNA oligos that permit specific binding, no evidence of condensate formation was observed, such as the accumulation of DNA in the well. We speculate that this may be due to the short length of the individual DNA fragments used in the assay.

To test whether increasing DNA length enables the formation of YY1–DNA assemblies beyond simple one-to-one binding complexes, we employed a single-molecule DNA curtain assay using λ-phage DNA (∼48.5 kb) (Fig. 1C)^26^. In this assay, hundreds of long DNA molecules are aligned on a lipid bilayer and extended by buffer flow against nanofabricated barriers, enabling high-throughput observation of protein–DNA interactions at the single-molecule level. DNA curtain assays have been widely used to investigate protein-induced DNA compaction and condensation at the single-molecule level^27–31^. In the current study, YY1 was fluorescently labeled with quantum dots (Q.dot) and introduced into flowcells containing single-tethered DNA curtains. At both 1 nM and 5 nM YY1, discrete fluorescent puncta were detected along the DNA (Fig. 1D). Strikingly, at 5 nM, we observed larger assemblies stably associated with the DNA. Quantitative analysis of fluorescence intensity revealed that these assemblies correspond to multimeric YY1 complexes, with detectable binding beginning at the dimer level, assuming the signals observed at 1 nM represent monomeric YY1 binding events. Because the labeling efficiency of YY1 is lower than 100%, these estimates likely underestimate the true degree of multimerization. These results demonstrate that even at low nanomolar concentrations, YY1 can form multimeric complexes on long DNA substrates, suggesting that the DNA length enables cooperative interactions that are not captured in short-probe assays.

To define the sequence specificity of YY1 on long DNA substrates, we mapped the positions of Q.dot-labeled signals along λDNA and compared them with predicted consensus YY1 motifs identified by MEME Suite analysis^32^ (Fig. 1E). Peaks in YY1 fluorescence intensity closely coincided with the predicted binding sites, confirming that YY1 retains sequence-specific recognition even within the extended and heterogeneous sequence context of λDNA. Notably, YY1 signals in GC-rich regions were more persistent, suggesting that local DNA flexibility, governed by base composition, can modulate the stability of YY1–DNA complexes^33,34^. To further examine whether multimeric YY1 assemblies exhibit a distinct positional preference, we reanalyzed the data by selectively extracting signals corresponding to putative multimeric YY1 species and reconstructed the binding profile (Supplementary Figure 1). The distribution of these multimeric signals closely resembled the overall binding pattern shown in Fig. 1E, indicating that multimeric YY1 retains a similar sequence-dependent binding tendency. These findings indicate that YY1 engages both high-affinity consensus motifs and nearby non-consensus sequences, and that its binding behavior is shaped by local sequence context. This context-dependent binding could facilitate the cooperative assembly of higher-order DNA–protein structures, setting the stage for the condensate formation phenomena described below.

### Concentration-dependent condensate formation by YY1

We next investigated whether varying the concentration of YY1 influences the formation of distinct higher-order structural assemblies on long DNA molecules. Using flow-assisted (0.2 mL/min) single-tethered DNA curtain assays, we found that at 10 nM YY1, the majority of λDNA molecules (81.7%) adopted bridged conformations, which occurred with nearly equal frequencies in two distinct configurations: intra-DNA bridging, in which condensate formation occurs within a single DNA molecule starting from its free ends, and inter-DNA bridging, in which assemblies form bridges between adjacent DNA molecules (Fig. 2A). Mapping the positions of bound YY1 revealed that inter-DNA bridging (trans) is mediated primarily through binding at internal DNA sites which lie in a region of high G/C content, whereas intra-DNA bridging (cis) originates from binding at terminal sites near the free ends of the DNA molecules (Fig. 2B). This preference for terminal regions may reflect the greater flexibility of A/T-rich DNA at the λDNA ends, which are more prone to collapsing, thereby facilitating single-DNA condensate formation. These observations suggest that even at relatively low nanomolar concentrations, YY1 can engage multiple binding configurations to organize DNA into alternative architectural states, potentially representing early intermediates in condensate formation.

**Figure 2:**
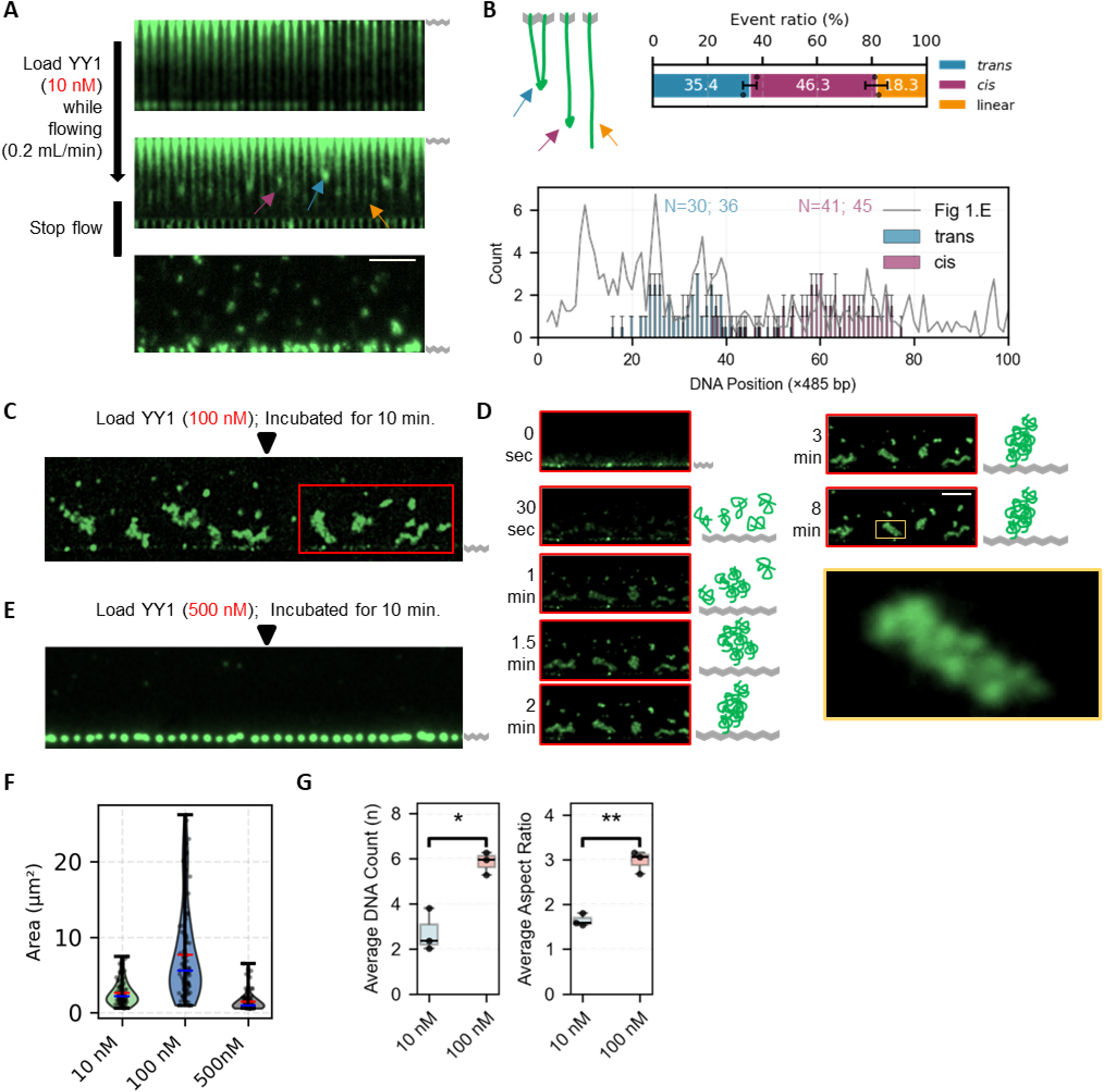
High concentrations of YY1-induces DNA condensate formation through multivalent crosslinking. **(A)** DNA curtain imaging of YY1-induced bridging and condensate formation. Top: aligned λDNA molecules under buffer flow. Middle: upon addition of 10 nM YY1, DNA molecules are drawn together from their free ends, forming bridging structures. Bottom: after stopping the flow, stable DNA condensates are observed, including trans-bridged and cis-condensed conformations. **(B)** Quantification of DNA bridging and condensate formation. Top: fraction of each structural class. Bottom: positional distribution of trans- and cis-interactions along DNA. Data are shown as mean ± SEM. **(C, E)** YY1 concentration-dependent DNA condensate formation. Representative images at (C) 100 nM and (E) 500 nM show extensive condensate formation on the flow cell surface. **(D)** Time-resolved condensate formation. Time-lapse images of the region indicated in (C) (red box) show progressive assembly of DNA condensates. Magnified views reveal diverse morphologies, ranging from extended linear assemblies to compact condensates. Cartoon schematics (right) illustrate representative conformations. The enlarged view (yellow box) highlights a densely packed, irregular condensate. Scale bars: 5 μm. **(F)** Size distribution of DNA condensates. Violin plots show condensate areas at 10 nM (green), 100 nM (blue), and 500 nM (orange). Red lines indicate the mean; blue lines indicate the median. **(G)** Quantification of condensate composition and morphology. The average number of DNA molecules per condensate increased from 2.7 ± 0.5 (10 nM) to 5.8 ± 0.3 (100 nM), accompanied by an increase in aspect ratio from 1.6 ± 0.1 to 3.0 ± 0.1. Statistical significance was assessed across three independent experiments (*P < 0.05, **P < 0.01).

After stopping the buffer flow, DNA molecules underwent spontaneous condensation. At 10 nM YY1, this process typically produced compact globular condensates derived from one or only a small number of DNA molecules (Bottom image of Fig. 2A). To our knowledge, no reliable measurements of the absolute nuclear concentration of YY1 have been reported in the literature. However, estimates of protein abundance indicate that YY1 falls within the top 25% of nuclear proteins, with copy numbers on the order of 1×10⁴ to 1×10⁵ molecules per cell^35^ . Assuming a nuclear volume of approximately 500 fL, this corresponds to an estimated YY1 concentration of roughly 100–500 nM, depending on cell types and cell cycle stages. Strikingly, when the YY1 concentration was increased to 100 nM, we unexpectedly observed the formation of much larger condensates in which multiple DNA molecules were linked together (Fig. 2C, 2D; Supplementary Movie 1). These assemblies were dynamic and loosely associated, indicating a different mode of organization from that seen at 10 nM (Fig. 2A, Supplementary Movie 2). The appearance of such multi-DNA condensates at intermediate concentration was remarkable, as it suggests a concentration window in which YY1 promotes a distinct, less rigid condensate state rather than simply driving progressive compaction. At 500 nM YY1, by contrast, the condensates consisted of multiple DNA molecules strongly linked into small, highly compact, and mechanically rigid structures (Fig. 2E), a condition in which the DNA also tended to adhere non-specifically to the DNA-curtain barrier surface. These results reveal that YY1 concentration not only modulates condensate size and compactness but also determines the number of DNA molecules incorporated and the physical nature of inter-DNA connections.

To quantitatively characterize YY1-dependent condensate formation, we analyzed condensates generated at 10 nM and 100 nM YY1 using area measurements based on Gaussian mixture modeling^36,37^ (Fig. 2F), where “area” refers to the pixel region over which the fluorescence signal spreads in the images. At 10 nM (n = 31 condensates), most condensates appeared as small assemblies in which one or only a few DNA molecules were confined within a single fluorescent punctum, whereas at 100 nM (n = 12), they more frequently comprised larger structures formed by the loose association of multiple DNA molecules into extended assemblies. Although there were clear differences in condensate area between the 10 nM and 100 nM conditions, condensates induced at 500 nM did not form larger structures due to aggregation and instead exhibited areas comparable to those at 10 nM. Morphologically, 10 nM condensates were generally smaller and nearly circular, while the 100 nM linked multi-DNA condensates were larger and exhibited significantly higher aspect ratios (Fig. 2G). These analyses reinforce the idea that the intermediate concentration state (100 nM), less rigid condensate architecture with unique assembly kinetics, is not a simple midpoint between low- and high-concentration.

### Contrasting solid-like DNA scaffolds and liquid-like YY1 dynamics within condensates

Given that the DNA condensates observed at intermediate (100 nM) YY1 concentrations displayed aspect ratios that deviated markedly from the spherical morphology typical of droplets formed as a result of LLPS, we next examined the internal dynamics of DNA and YY1 within these structures^38,39^. YY1 was labeled with Q.dot to enable real-time visualization, with the labeling density kept sufficiently low so that not all YY1 molecules were tagged. This sparse labeling allowed individual Q.dot-labeled YY1 molecules to be distinguished, revealing that Q.dot signals localized within the condensates (Supplementary Movie 3 and 4). Unexpectedly, YY1 molecules within these assemblies exhibited pronounced dynamic mobility, whereas the associated DNA scaffolds remained immobile (Fig. 3A).

**Figure 3:**
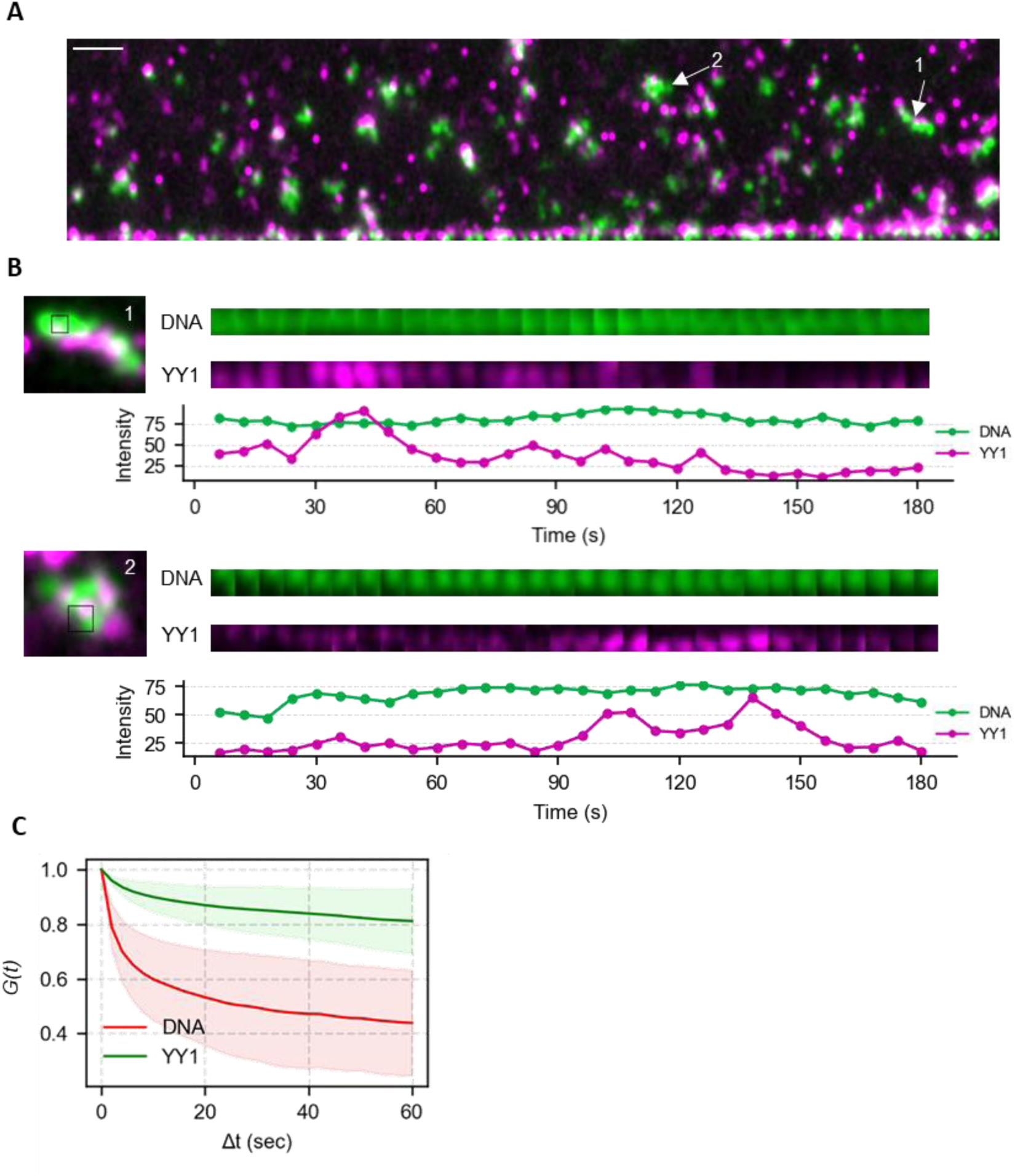
Distinct physical properties of YY1 and DNA within YY1–DNA co-condensates. **(A)** Representative fluorescence image of quantum dot–labeled YY1 (magenta) colocalized with DNA condensates (green). Scale bar, 5 μm. Time-lapse snapshots (below) show particle tracking of YY1 and DNA within a condensate. **(B)** Time-resolved analysis of DNA and YY1 fluorescence signals within single condensates. Representative kymographs of DNA (green) and Q.dot-labeled YY1 (magenta) intensities extracted from fixed regions (1 and 2, indicated in panel A) within individual condensates are shown. The corresponding fluorescence intensity traces over time are plotted below each kymograph. **(C)** Autocorrelation functions of fluorescence intensity fluctuations for YY1 (magenta) and DNA (green), averaged over 45 condensates.

To quantitatively characterize these dynamics, we tracked the fluorescence intensities of DNA and individual Q.dot-labeled YY1 molecules within fixed regions of a single condensate over time (Fig. 3B). While the DNA signal remained largely constant, indicating a stable and immobile scaffold, the YY1 signal exhibited pronounced temporal fluctuations, repeatedly increasing and decreasing in intensity. Importantly, even during intensity decreases, the YY1 signal did not fall to background levels, arguing against Q.dot blinking as the primary source of these fluctuations. Notably, YY1 was loaded under continuous buffer flow, ensuring that unbound molecules were efficiently washed away and that only DNA-associated YY1 molecules remained within the field of view. Despite this, the observed intensity fluctuations persisted, indicating that YY1 molecules dynamically redistribute within a single condensate rather than exchanging with the bulk solution. Although the analysis was performed within a spatially fixed region of a single condensate, the YY1 signal intermittently appeared and disappeared within this region, and even within the same condensate, regions completely lacking detectable YY1 signal were observed. This spatial heterogeneity suggests that YY1 molecules preferentially accumulate in regions where YY1 is already present, indicative of cooperative binding. Together, these observations reveal that, despite the solid-like nature of the DNA scaffold, YY1 molecules exhibit highly dynamic and cooperative rearrangements within individual condensates.

To further quantify the temporal stability of condensate structures, we computed the auto-correlation function *G*(Δ*t*), defined as the spatial average of the normalized pixel-wise product between image frames separated by a time lag Δ*t* (Fig. 3C; See Methods). As shown in Fig. 3C, the auto-correlation of the DNA signal (green) decayed more slowly and remained at relatively high values over the entire time window, indicating that the spatial distribution of DNA within condensates is temporally stable. In contrast, the auto-correlation of the YY1 signal (red) decreased more rapidly, reflecting faster rearrangement or turnover of the protein component within the same structures. The clear separation between the two curves again suggests that, while the DNA scaffold maintains structural persistence, YY1 molecules exhibit greater dynamic exchange or internal mobility within the condensates. Together, these results indicate that YY1-DNA co-condensation produces mechanically stable condensates scaffolded by DNA, inside which YY1 remains highly mobile.

### YY1 concentration-dependent mechanical stability of YY1-DNA co-condensates

To assess the stability and reversibility of YY1-induced DNA condensates, we applied increasing buffer flow rates to disassemble pre-formed structures. Condensates were first generated by incubating λDNA with 10 nM, 100 nM, or 500 nM YY1 without flow and then subjected to a 0.5 mL/min buffer flow for 10 minutes. Most condensates formed at 10 nM YY1 rapidly transitioned into extended forms, often resolving into linear DNA that retained discrete, localized puncta along its length (Fig. 4A, top). These puncta remained stable over time and were distributed predominantly along the DNA molecule. At 100 nM, weakly linked multi-DNA condensates displayed dynamic structural rearrangements before stabilizing either as extended linear forms or as clearly defined bridging structures connecting multiple DNA molecules through inter-DNA contacts (Fig. 4A, middle). These behaviors suggest that inter-DNA linkages in the weakly linked state can reorganize under shear into alternative architectural states. At the highest concentration (500 nM), strongly linked multi-DNA condensates were far more resistant to flow-induced rearrangement (Fig. 4A, bottom).

**Figure 4:**
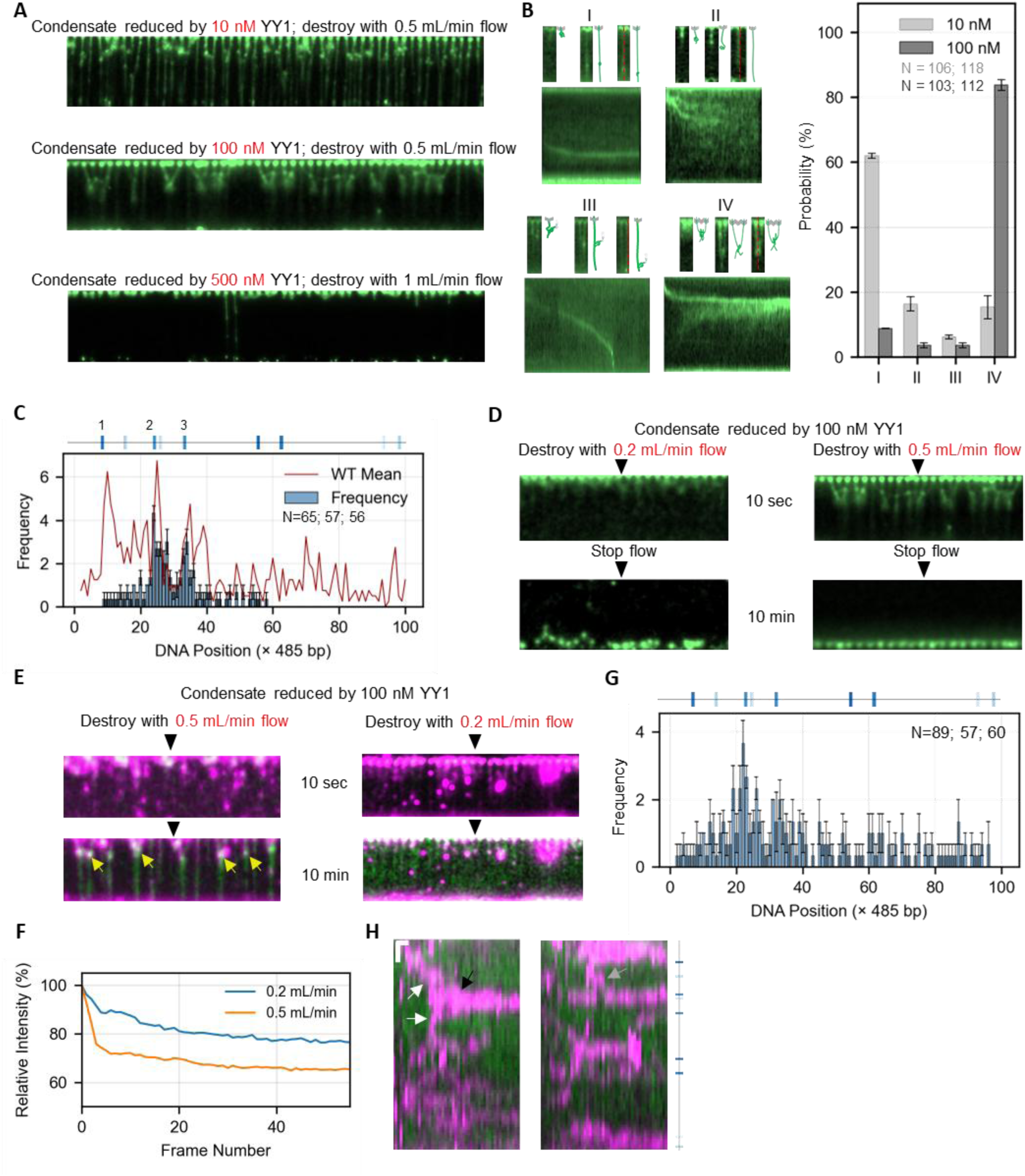
Force-dependent disassembly and motif-anchored reorganization of YY1–DNA condensates. **(A)** Final steady-state structures of YY1-induced DNA condensates after exposure to a strong buffer flow (0.5 mL/min), shown for increasing YY1 concentrations (10, 100, and 500 nM; top to bottom). **(B)** Representative kymographs showing the response of condensates formed at 100 nM YY1 to high flow (0.5 mL/min). The bar graph (right) summarizes the relative frequencies of these four disassembly modes under 10 nM and 100 nM YY1 conditions (mean ± SEM; sample sizes indicated). **(C)** Spatial distribution of residual foci along λDNA after condensate disruption (mean ± SEM). The red line indicates the distribution of YY1 binding events along WT λDNA (Fig. 1E). **(D)** Flow-dependent reversibility of condensates formed at 100 nM YY1. Top: condensate disruption under moderate (0.2 mL/min) and high (0.5 mL/min) flow. Bottom: states immediately after flow cessation. **(E)** Representative time-lapse images of single quantum dot–labeled YY1 molecules on λDNA under moderate (0.2 mL/min, left) and high (0.5 mL/min, right) flow. Yellow arrows indicate YY1 binding to DNA. **(F)** Time-dependent decay of quantum dot fluorescence signals under 0.2 and 0.5 mL/min flows. **(G)** Histogram of YY1 positions remaining on λDNA after condensate disruption (mean ± SEM). **(H)** Kymographs of representative fluorescence images of YY1–DNA condensates under moderate flow (0.2 mL/min). (Left) fusion of two diffusing YY1 molecules (white arrows) into a stable oligomer (black arrow) near a binding site, and (Right) transient dissociation and re-binding of YY1 at a specific site (gray arrow).

Closer inspection of the events revealed two main modes of disassembly dynamics for condensates formed at 10 nM YY1: most often, DNA extended into a linear form while retaining localized puncta (I, 62% at 10 nM), whereas complete linearization without puncta (II, 16%) was relatively rare (Fig. 4B). For condensates formed at 100 nM YY1, the more common behavior was the formation of bridging structures between adjacent DNA molecules via puncta (IV, 83.8% at 100 nM), while a less frequent mode involved the unraveling of one DNA molecule, with the others sliding along the flow direction while remaining aggregated, before ultimately dissociating (III, 3.7%) (Fig. 4B). In all cases, the residual puncta persisted over time. Mapping these stable puncta to the λDNA sequence revealed a high overlap with predicted YY1 binding sites (Fig. 4C), indicating that these persistent substructures are anchored by sequence-specific YY1–DNA interactions. Among the three YY1 consensus motifs near the barrier, the motif closest to the barrier (motif 1) is rarely used during cross-structure formation, whereas motifs 2 and 3 are preferentially engaged. This preference likely reflects geometric constraints: using motif 1 would require an unrealistically sharp DNA bending angle, while motifs 2 and 3 allow cross-bridging with a much more favorable, gentler bend. Together, these results suggest that YY1-induced DNA condensates are stabilized by a combination of strong, sequence-specific contacts and weaker, non-specific interactions, with the specific components providing mechanical stability that resists complete disassembly.

Importantly, once YY1-induced DNA condensates were disassembled by buffer flow (0.5 mL/min), neither condensates formed at 10 nM nor condensates formed at 100 nM reverted to their original compacted states, even after the flow was stopped again, indicating that the disassembly process is irreversible under these conditions. At intermediate flow rates (0.2 mL/min), weakly linked multi-DNA condensates formed at 100 nM YY1 did not fully disassemble; instead, they underwent collective deformation that followed the contour of the nanofabricated barrier (Fig. 4D). The persistence of discrete, particle-like DNA substructures and inter-DNA linkages under mild shear stress indicates that weakly linked condensates can undergo reversible mechanical rearrangements without complete disassembly. Together, these results show that YY1-induced DNA condensates exhibit tunable mechanical stability as a function of YY1 concentration and the sequence-specific binding sites play a key role in anchoring residual structures.

### YY1-DNA co-condensate stabilization by non-specific and specific YY1 binding

To investigate the molecular basis of condensate formation by YY1, we tracked the behavior of individual YY1 molecules using Q.dot-labeled protein under flow conditions. Weakly linked multi-DNA condensates were first assembled at 100 nM YY1, and then a buffer flow of 0.5 mL/min was applied to induce partial dissociation of YY1. Under these conditions, most Q.dot-labeled YY1 signals rapidly disappeared from the DNA, indicating the loss of loosely bound molecules (Fig. 4E, left & 4F). In contrast, the remaining YY1 molecules were consistently localized at discrete puncta within the condensates (Fig. 4E, left), suggesting that these sites act as structural anchors stabilized by sequence-specific YY1–DNA interactions (Fig. 4G).

Under weaker flow conditions that led to irreversible disassembly (0.2 mL/min), we observed numerous Q.dot-labeled YY1 molecules diffusing along λDNA (Fig. 4E, right & 4H). These molecules frequently underwent transient binding and dissociation at various positions, moving in an apparent one-dimensional random walk independent of the flow direction. During this diffusion, some YY1 molecules merged to form larger aggregates (Fig. 4H, left). We also observed events in which YY1 molecules transiently stabilized at one specific site before relocating to another and returning to this site (Fig. 4H, right). Such mobility and transient interactions suggest that both non-sequence-specific and sequence-specific bindings alternately occur, reflecting the dynamic behaviours of YY1 along the DNA. Because applying a 0.5 mL/min flow removes nearly all non-specifically bound YY1 molecules—and, under these conditions, the condensates can no longer reform once they are disrupted (Fig. 4D, right) —it becomes clear that these non-specific interactions are essential for the initial stages of condensate formation and assembly.

In sum, YY1-induced DNA condensation is initiated by a population of dynamically interacting, non-specifically bound molecules that bring distant DNA regions into proximity, in which specific binding sites stabilize the resulting architecture. This conclusion contrasts with earlier models suggesting that YY1 bridges pre-defined promoter and enhancer regions by binding to specific sites^40^, although it remains possible that both mechanisms operate depending on the chromatin context and YY1 concentration.

### YY1-subregion-specific contributions to condensate formation

To dissect the molecular requirements for YY1-induced DNA condensation, we generated a series of subregion-deletion mutants targeting the IDR and DNA-binding domain of YY1. Specifically, we engineered five variants: YY1-ΔE/D (glutamate/aspartate-rich region deletion), YY1-ΔH (histidine-rich region deletion), YY1-ΔG/K (glycine/lysine-rich region deletion), YY1-ΔIDR (full IDR deletion), and YY1-ΔZF (zinc finger domain deletion) (Fig. 5A). Using EMSA, we confirmed that all mutants except YY1-ΔZF retained sequence-specific DNA-binding activity at low concentrations (Fig. 5B, Supplementary Figure 2A).

**Figure 5:**
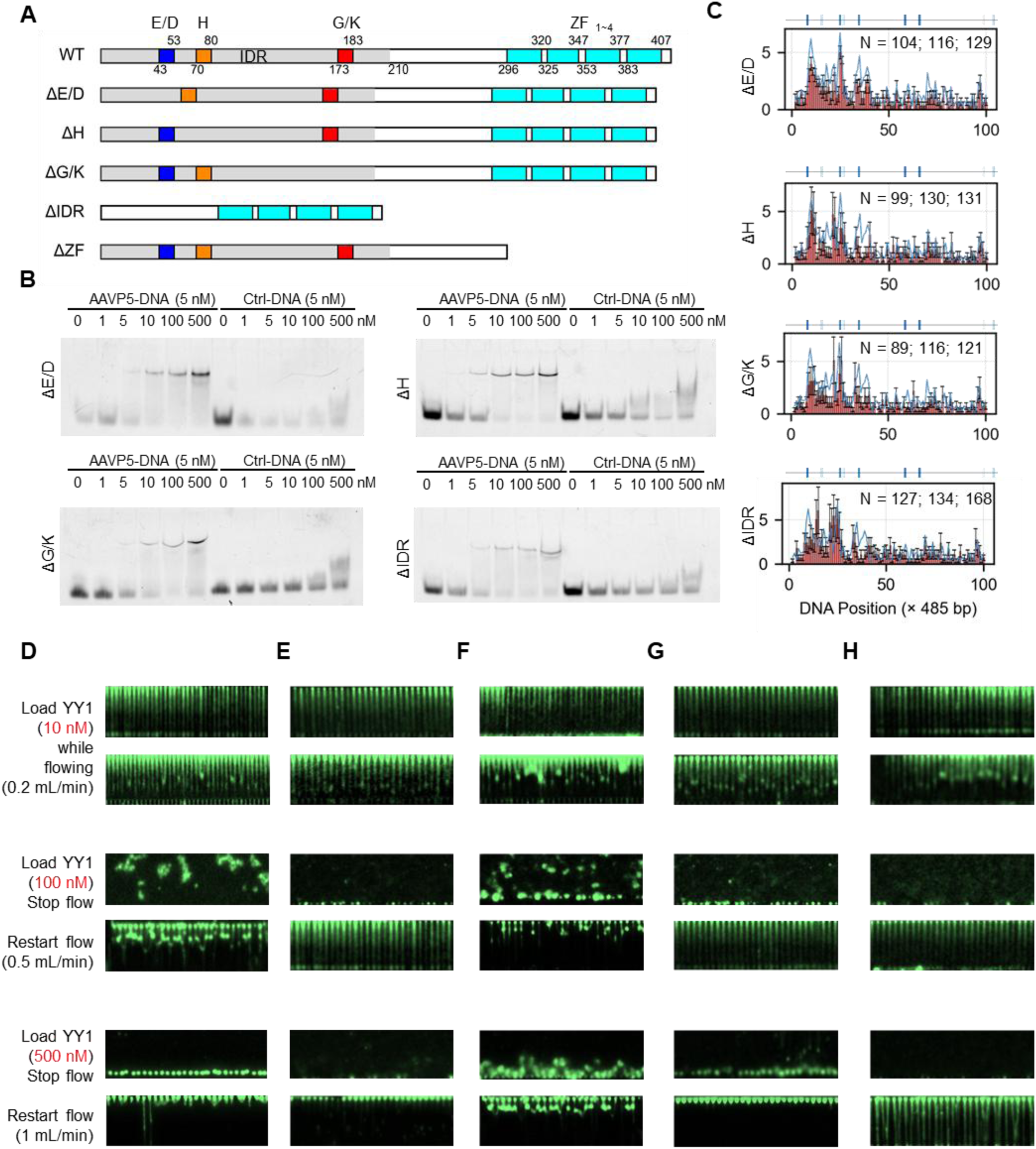
The G/K region and zinc finger domains are required for YY1-induced DNA condensation and stability. **(A)** Domain organization of wild-type YY1 and deletion mutants: ΔE/D (acidic region), ΔH (histidine-rich region), ΔG/K (glycine/lysine-rich region), ΔIDR (entire intrinsically disordered region), and ΔZF (zinc fingers). **(B)** Electrophoretic mobility shift assay (EMSA) of DNA binding by each YY1 variant using AAVP5 promoter DNA or control DNA. DNA concentration was fixed at 5 nM, with YY1:DNA molar ratios of 0, 0.2, 1, 2, 20, and 100. **(C)** Binding profiles derived from DNA curtain assays for each YY1 variant (mean ± SEM). **(D–H)** DNA condensation phenotypes of YY1 variants at increasing concentrations. Representative images show DNA bridging at 10 nM, soft multi-DNA condensation at 100 nM, and hard multi-DNA condensation at 500 nM.

**Figure 6:**
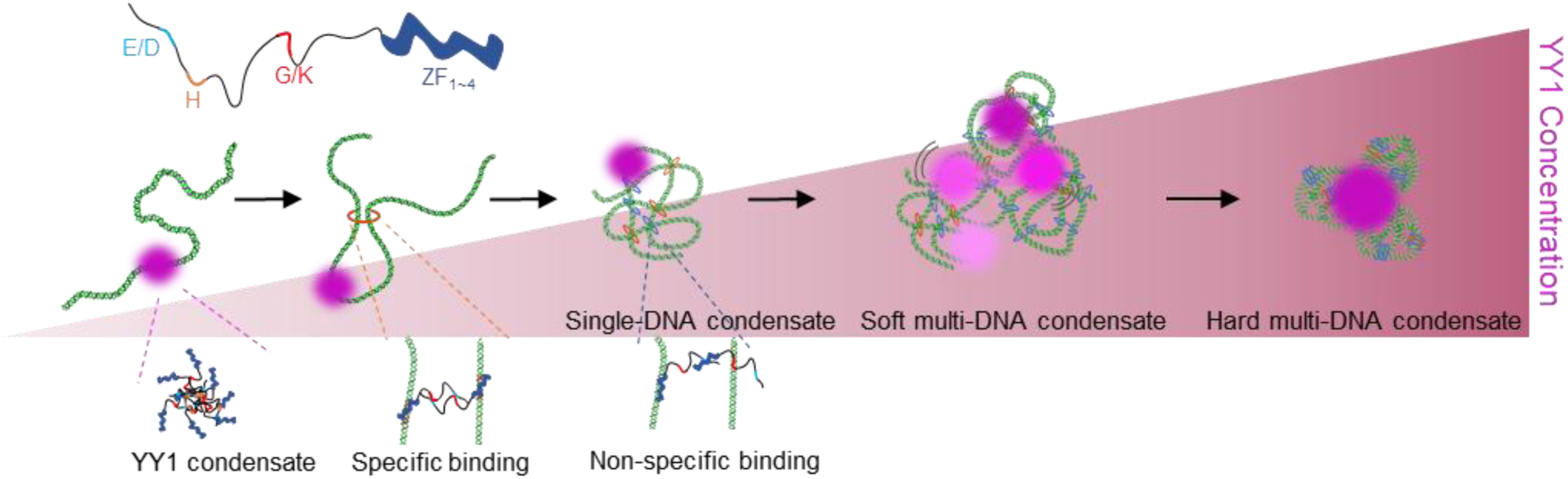
Schematic model illustrating how increasing YY1 concentration drives the transition from individual DNA-bound YY1 to single-DNA condensate, weakly-linked (soft) multi-DNA condensates and ultimately to strongly-crosslinked (hard) multi-DNA condensates. These assemblies are mediated by a combination of sequence-specific binding through the zinc finger (ZF) domain and multivalent, non-specific interactions involving intrinsically disordered regions (IDRs) and residue regions (e.g., E/D and G/K regions), collectively promoting higher-order DNA organization.

To determine the role of each YY1 domain in DNA condensate formation, we next performed DNA curtain assays using these mutants (Fig. 5C). As the first condition, to examine whether low concentrations (10 nM) of YY1 mutants induce compaction of DNA molecules, we loaded 10 nM YY1 (WT or each mutant) onto DNA under a slow buffer flow of 0.2 mL/min and observed the molecules without stopping the flow (Top two rows of Fig. 5D to H). As a result, we found that none of the mutants had lost the ability to compact DNA. Interestingly, YY1-ΔIDR still retained the ability to compact DNA. Given that the spacer region from residue 210 to 295 contains only a limited number of positively charged amino acids (Fig. 1A) and is therefore unlikely to bind DNA, this compaction is likely caused by the four zinc finger domains bridging distant DNA segments. Consistent with this interpretation, the ΔZF mutant showed neither detectable DNA binding nor compaction under the same conditions (Supplementary Figure 2B), further supporting the idea that the zinc finger domains are essential for mediating DNA bridging and compaction.

As the third condition, we loaded high concentrations (500 nM) of YY1 onto the DNA curtain, stopped the buffer flow, incubated for 10 minutes, and then resumed the buffer flow at 0.5 mL/min (Bottom two rows of Fig. 5D to H). Wild-type YY1 formed hard multi-DNA condensates (condensates that are scarcely disrupted even under a buffer flow of 1.0 mL/min) during the 10-minute incubation. Surprisingly, this hard multi-DNA condensate was also observed when the mutant lacking the IDR was loaded (Bottom two rows of Fig. 5E). This result suggests that hard multi-DNA condensates were formed through increased bridging between DNA segments by the four zinc finger domains that mediate DNA compaction. Interestingly, when the two positively charged regions—the H-rich region and the G/K-rich region—are present (YY1-ΔE/D), soft multi-DNA condensates were formed (condensates that are partially disrupted under a buffer flow of 1.0 mL/min) (Bottom two rows of Fig. 5F). These results again suggest that the H-rich region and the G/K-rich region play important roles in the formation of soft multi-DNA condensates.

At the same time, the fact that the deletion of E/D-rich region impairs the formation of hard multi-DNA condensates, yet YY1-ΔIDR does not, indicates that the E/D-rich region likely functions in concert with other IDR subregions to form hard multi-DNA condensate. Importantly, the observation that complete removal of the IDR does not impair, but rather permits robust hard multi-DNA condensate formation strongly supports a model in which the IDR exerts an overall autoinhibitory effect on zinc finger-mediated DNA binding and bridging^41^. In contrast, when the G/K-rich region and the E/D-rich region are present (YY1-ΔH), soft multi-DNA condensates were not formed, and hard DNA condensates were formed instead (Bottom two rows of Fig. 5G). This result suggests that the H-rich region plays a negligible role in forming hard multi-DNA condensates. Also, when the H-rich region and the E/D-rich region were present (YY1-ΔG/K), neither soft multi-DNA condensates nor hard multi-DNA condensates were formed (Bottom two rows of Fig. 5H). This is notable because again, YY1-ΔIDR does not impair hard multi-DNA condensate formation, whereas deleting only the G/K-rich region does. This contrast suggests that the G/K-rich region also does not act alone but instead cooperates with other IDR subregions to promote the formation of hard multi-DNA condensates. A plausible model is that the negatively charged E/D-rich region and the positively charged G/K-rich region engage in intramolecular or intermolecular charge-complementary interactions, which enhance the efficiency of zinc finger-domain-mediated DNA bridging, thereby promoting the formation of mechanically robust hard multi-DNA condensates.

In summary, among the characteristic regions within the IDR of YY1, the two positively charged regions contribute to the formation of soft multi-DNA condensates, and the E/D-rich region appears to engage in charge-complementary interactions with the G/K-rich region, enhancing zinc finger-domain-mediated DNA bridging and thereby promoting the assembly of hard multi-DNA condensates.

## DISCUSSION

Our study reveals that YY1 engages in highly modular and concentration-dependent interactions with long double-stranded DNA, leading to the formation of distinct condensate types through the combined actions of its DNA-binding zinc finger domains and IDR. At moderately high concentrations, YY1 assembles into “soft” multi-DNA condensates, within which YY1 molecules exhibit rapid, liquid-like mobility, whereas the associated DNA remains relatively immobile and behaves in a solid-like manner. Notably, these soft multi-DNA condensates were formed through the combined action of specifically bound YY1 molecules anchored at consensus motifs and non-specifically bound YY1 molecules. This mode of condensation critically depends on the histidine-rich and glycine/lysine-rich regions within the IDR.

At high concentrations, zinc finger domain–mediated bridging dominates, leading to robust “hard” condensates. Importantly, the fact that mutations in distinct YY1 domains selectively disrupt either hard or soft multi-DNA condensate formation demonstrates that these two condensate types are generated by fundamentally different mechanisms. Together, these findings challenge the traditional view that higher YY1 concentrations merely increase occupancy at specific DNA motifs. Instead, they reveal that YY1 can shift the material state of DNA–protein assemblies in a concentration-dependent manner, suggesting that the physical properties of YY1–DNA co-condensates themselves constitute an additional regulatory layer with the potential to shape downstream transcriptional outcomes.

Previous studies have reported that high concentrations of YY1 can form droplets via LLPS, and that such protein condensates are driven primarily by the H-rich region within its IDR^20^. Consistent with this, our domain-deletion analysis confirmed that removal of the H-rich region abolished the formation of soft multi-DNA condensates. However, we also found that deletion of the G/K-rich region, in addition to the H-rich region, similarly prevented soft condensate formation. This finding reveals that the interactions necessary for soft multi-DNA condensate assembly are not solely those that underlie protein–protein condensation in LLPS, but also involve additional interaction modes—likely multivalent electrostatic contacts between the positively charged IDR subregions and DNA. Thus, while the H-rich region is a well-established driver of YY1 protein condensates^20^, our results demonstrate that the G/K-rich region provides an equally essential contribution in the DNA context, expanding the conceptual framework for how YY1’s IDR mediates distinct condensate types.

Muzzopappa et al. demonstrated that the material properties of DNA-based condensates are strongly governed by DNA length, with short DNA fragments preferentially forming liquid-like assemblies and longer DNA molecules transitioning toward gel- or solid-like state^42^. Their work established an intrinsic link between polymer length and condensate fluidity, suggesting that the physical properties of DNA itself can dictate whether a condensate behaves as a dynamic droplet or a more rigid aggregate. In contrast to this length-dependent paradigm, our study of YY1-induces DNA condensation reveals a qualitatively different layer of regulation: within YY1-DNA co-condensates, YY1 retains liquid-like dynamics, whereas the DNA scaffold exhibits solid-like or highly constrained behavior. Thus, rather than DNA length alone determining condensate material properties, our findings indicate that protein–DNA compositional asymmetry can decouple the dynamics of components within the same condensate.

The formation of soft multi-DNA condensates may have important physiological implications for YY1’s function in gene expression regulation. In our model, multivalent non-specific binding of YY1 to DNA promotes local condensation of chromatin segments, thereby increasing the likelihood of bridging between distant specific binding sites. Such a mechanism could facilitate YY1’s established role in mediating enhancer–promoter communication, where transient, reversible DNA condensate formation would promote the juxtaposition of regulatory elements without imposing rigid architectural constraints. The dynamic, liquid-like behavior of YY1 within these soft multi-DNA condensates may further enable rapid reconfiguration of DNA–protein contacts in response to transcriptional cues, allowing YY1 to efficiently sample and connect its consensus motifs across the genome^43^.

In contrast, hard multi-DNA condensates may serve a fundamentally different biological role if any. Their formation is driven predominantly by zinc finger-domain-mediated DNA bridging, resulting in mechanically rigid architectures that resist disruption even under strong shear. Such stable DNA condensation could contribute to long-term chromatin compaction, gene silencing, or the maintenance of higher-order chromatin domains in contexts where structural integrity is prioritized over dynamic regulatory exchange. Unlike soft multi-DNA condensates, which allow rapid rearrangement of DNA–protein interactions, hard multi-DNA condensates are likely to restrict DNA accessibility and limit the mobility of associated factors, thereby establishing durable repressive states. The ability of YY1 to form either soft or hard multi-DNA condensates suggests that the protein may switch between dynamic regulatory scaffolds and stable architectural anchors in a context-dependent manner.

In this study, we demonstrated that the relative abundance of soft and hard multi-DNA condensates can be drastically altered by changing the concentration of YY1; however, in the cellular environment, these two condensate types are likely to coexist. We propose that the ratio between soft and hard multi-DNA condensates could modulate the accessibility of transcriptionally essential factors—such as the Mediator complex and other transcription factors—to DNA condensates, thereby influencing transcriptional regulation. This represents a model distinct from a simple monotonic relationship in which transcription output increases proportionally with transcription factor abundance, instead suggesting a more nuanced regulatory mechanism governed by condensate composition.

A limitation of this study is that all mechanistic analyses of condensate formation were performed *in vitro* using reconstituted systems, which do not fully capture the complexity of the nuclear environment, such as chromatin compaction, nucleosome positioning, and the presence of additional cofactors. Moreover, the physiological concentrations of YY1 and the precise distribution of its post-translational modifications in different cell types remain to be determined. Future *in vivo* experiments, including live-cell imaging and targeted mutagenesis, will be essential to validate the soft/hard multi-DNA condensate model under physiological conditions. Despite these limitations, our findings establish a domain-level framework in which YY1 can form two distinct types of DNA condensates through separable structural modules, providing a new conceptual basis for how transcription factors integrate specific and non-specific DNA interactions to regulate genome organization and transcriptional output.

## METHODS

### Protein expression and purification of YY1 and its mutants

The coding sequences for human YY1 and its mutants were codon-optimized for *Escherichia coli* expression and synthesized by Eurofins Genomics. An N-terminal His₆–HA₂ tag was added for affinity purification. Each construct was cloned into the pET-28a(+) expression vector (Novagen) using the NheI and XhoI restriction sites, and the resulting plasmids were transformed into *E. coli* DH5α for amplification. Plasmids were isolated using the Wizard Plus Midipreps DNA Purification System (Promega). For protein expression, the verified plasmids were transformed into *E. coli* BL21 (DE3) competent cells (New England Biolabs, C2527H). Cells were cultured in LB medium supplemented with 100 μg/mL ampicillin at 37°C and induced with 0.5 mM IPTG at an OD₆₀₀ of ∼0.6. After overnight expression at 16°C, cells were harvested by centrifugation and lysed by sonication in buffer containing 50 mM Tris-HCl (pH 7.5), 300 mM NaCl, 10 mM imidazole, and 1 mM DTT. Protein purification was carried out following the previously established procedures. Cell lysates were cleared by centrifugation and applied to a Ni–NTA agarose column (QIAGEN) pre-equilibrated with lysis buffer. Following washing with lysis buffer containing 20 mM imidazole, His-tagged YY1 proteins were eluted with 250 mM imidazole. Full-length YY1 and all mutants except ΔZF were further purified by DNA affinity chromatography, whereas YY1-ΔZF was purified by size-exclusion chromatography. Protein purity (>90%) was confirmed by SDS–polyacrylamide gel electrophoresis (PAGE), and protein concentrations were determined using the BSA assay.

### Electrophoretic mobility shift assay

To assess sequence-specific DNA binding by YY1 and its mutants, EMSA experiments were performed using native PAGE. A 20-bp oligonucleotide containing the YY1 consensus binding site from the adeno-associated virus P5 promoter (AAVP5-DNA: 5′-CGTTTCAAAATGGAGACCCT-3′) was used as the target, and a randomized oligonucleotide of identical length (Ctrl-DNA: 5′-TTAGCTATGAACTGAGTCGTCATCT-3′) served as a negative control. The complementary strand of each oligonucleotide was labeled at the 5′ end with Alexa Fluor 546N for fluorescence detection. Protein–DNA binding reactions were prepared in 10 µL volumes containing binding buffer (10 mM Tris-HCl, pH 7.5; 50 mM NaCl; 1 mM MgCl₂; 0.1 mg/mL BSA), 5 nM fluorescently labeled DNA, and increasing concentrations of YY1 (0–500 nM). After incubation for 20 minutes at room temperature, samples were loaded onto a 5–20% native PAGE gel (e-PAGEL HR, ATTO; Cat# 2331970) and electrophoresed at 4°C in TBE buffer. Fluorescent signals were detected using an iBright FL1500 Imaging System (Thermo Fisher Scientific).

### DNA curtain assay

Single-tethered DNA curtains were prepared as previously described using λ-phage DNA^44^ (New England Biolabs). Flowcells were cleaned with water and equilibrated with 3 mL of lipid buffer (10 mM Tris-HCl, pH 7.5; 100 mM NaCl). A lipid mixture with a molar ratio of DOPC : PEG2000–DOPE : biotin–DOPE = 300 : 10 : 1 was injected into the flowcell in three separate 1-mL aliquots, with 5-minute incubations between injections. The flowcell was then rinsed with 3 mL of lipid buffer and incubated for 1 hour at room temperature to stabilize the lipid bilayer. Next, 3 mL of DNA-binding buffer (25 mM Tris-HCl, pH 7.5; 2 mM MgCl₂; 2 mM DTT; 0.2 mM ZnCl₂; 100 mM NaCl; 1.0 mg/mL BSA) was flushed through the chamber and incubated for 5 minutes. Streptavidin (1 mg/mL, New England Biolabs, N7021S) was then injected in two 400-μL aliquots, each followed by a 5-minute incubation. After washing with 3 mL of DNA-binding buffer, biotinylated λDNA (500 ng/mL) was introduced and incubated for 10 minutes to allow surface attachment via the biotin–streptavidin interaction. Following DNA immobilization, the flowcell was mounted on a fluorescence microscope. DNA was extended by applying a buffer flow of 0.3 mL/min for 5 minutes, followed by a 5-minute static incubation. To enhance DNA visualization, YOYO-1 intercalating dye (1 nM final concentration) was introduced at a flow rate of 0.2 mL/min for 10 minutes. Fluorescence imaging was performed using total internal reflection fluorescence microscopy, and images were acquired at multiple time points depending on the experimental setup.

### Image autocorrelation analysis

To quantify the temporal stability of fluorescence patterns, we computed the time-dependent autocorrelation function *G*(*t*), which measures the similarity between an image acquired at time *t*_0_ and a subsequent image at *t*_0_ + *t*. Each fluorescence image was treated as a two-dimensional matrix of pixel intensities *I*(*x*, *y*). For a given time point, pixel intensities were normalized to remove global intensity bias and enable comparison across time. Specifically, each pixel intensity was mean-centered and scaled by the standard deviation:

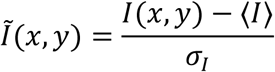

where ⟨*I*⟩ and *σ*_*I*_ denote the mean and standard deviation of pixel intensities within the image, respectively. To compute the autocorrelation at time lag *t*, the normalized images at *t*_0_ and *t*_0_ + *t*, denoted as 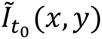 and 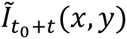, were multiplied element-wise to generate a correlation map:

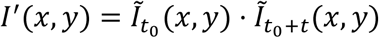

The autocorrelation function *G*(*t*) was then calculated as the spatial average over all pixels:

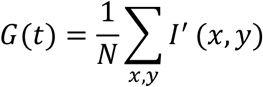

where *N* is the total number of pixels. A high value of *G*(*t*) indicates strong similarity between the two images, corresponding to stable or slowly evolving fluorescence patterns. In contrast, a low or near-zero value of *G*(*t*) reflects reduced similarity, indicative of dynamic rearrangements in the fluorescence signal over time.

## ACKNOWLEDGEMENT

We would like to thank members of the Theoretical Biophysics laboratory at Kyoto University for discussions and assistance throughout this work. This work was supported by the Grant-in-Aid for Transformative Research Areas of Japan Society for the Promotion of Science (24H00882 to T.T.), the grant from the Takeda Science Foundation (to T.T.), the grant from the Shimazu Science Foundation (to T.T.), the grant from the Yamada Science Foundation (to T.T.).

## FIGURES

**Supplementary Figure 1:**
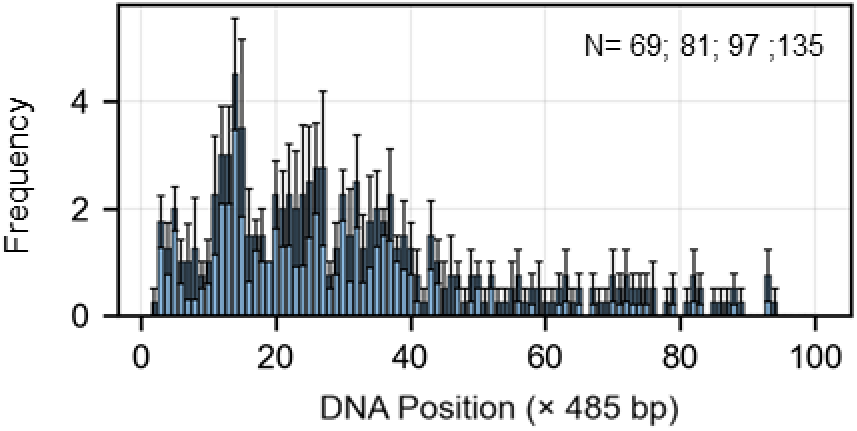
Histogram showing the YY1 binding events disruption (mean ± SEM) along DNA at 5 nM, calculated from fluorescence signals above the mean intensity threshold. Data are shown as mean ± SEM.

**Supplementary Figure 2:**
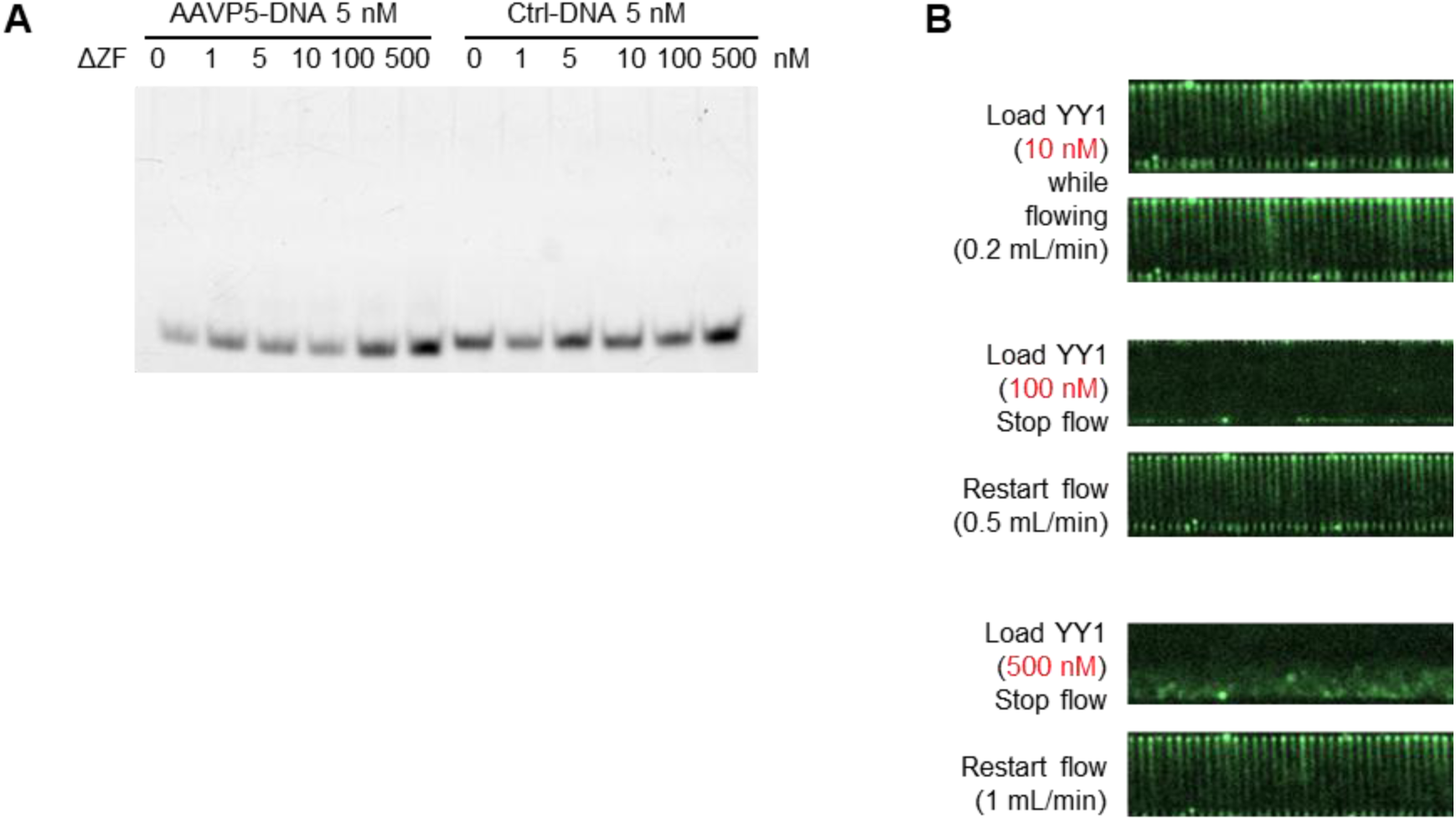
Loss of zinc finger domain impairs YY1 DNA binding and condensation. **(A)** Electrophoretic mobility shift assay (EMSA) of the ΔZF-YY1 variant using AAVP5 promoter DNA or control DNA. DNA concentration was fixed at 5 nM, with YY1:DNA molar ratios of 0, 0.2, 1, 2, 20, and 100. **(B)** DNA curtain images of ΔZF-YY1 at increasing concentrations (10, 100, and 500 nM). Unlike wild-type YY1, the ΔZF mutant shows markedly reduced DNA binding and fails to induce robust DNA condensation across all conditions.

